# Glucagon-like peptide-1 receptor C-terminal tail phosphorylation determines signalling responses from pancreatic β-cells

**DOI:** 10.1101/2025.11.01.686001

**Authors:** Yusman Manchanda, Natalie Wong, Annabelle Milner, Alejandra Tomas

## Abstract

The glucagon-like peptide-1 receptor (GLP-1R) is a class B G protein–coupled receptor that regulates glucose homeostasis by promoting cAMP production and insulin secretion in pancreatic β-cells. Although receptor phosphorylation is known to influence GLP-1R signalling, its functional relevance in β-cells remains incompletely understood. Here, we investigated the role of three C-terminal serine doublets (Ser441/442, Ser444/445, Ser451/452) in the human receptor using site-directed mutagenesis in INS-1 832/3 β-cells deleted for endogenous GLP-1R expression. GLP-1 stimulation induced serine phosphorylation of wild-type GLP-1R, which was abolished in all individual doublet as well as triple doublet phospho-null mutants. Basal phosphorylation was detected in wild-type receptors under unstimulated conditions, suggesting dynamic cycling between phosphorylated and unphosphorylated states. All mutant receptors trafficked normally to the plasma membrane, but the triple alanine doublet (AA) mutant exhibited reduced GLP-1-induced internalisation and significantly decreased plasma membrane diffusion under vehicle conditions, indicating altered receptor dynamics even in the absence of agonist. Importantly, the Ser451/452 doublet was required for GRK2 recruitment to the active GLP-1R. Functionally, the triple AA mutant showed impaired GLP-1–induced cAMP generation with minimal change in β-arrestin 2 recruitment, resulting in biased signalling. The cAMP defect was partially rescued by phosphodiesterase 4 (PDE4) inhibition with rolipram, implicating altered PDE engagement in the defective cAMP response. These findings demonstrate that both basal and agonist-induced GLP-1R phosphorylation are critical for receptor signalling in β-cells, revealing a previously unappreciated role for basal phosphorylation in shaping receptor behaviour and establishing a positive link between GLP-1R phosphorylation and β-cell signalling competence.

## Introduction

The glucagon-like peptide-1 receptor (GLP-1R) is a class B G protein–coupled receptor (GPCR) that plays a pivotal role in glucose homeostasis by mediating the insulinotropic and anorexigenic effects of incretin hormones (1). GLP-1R activation by endogenous ligands or pharmacological agonists stimulates adenylate cyclase (AC) through Gαs coupling, leading to elevations in intracellular cAMP and enhanced insulin secretion from pancreatic β-cells (2). Beyond classical G protein signalling, GLP-1R function is tightly regulated by post-translational modifications (PTMs), with receptor phosphorylation being particularly relevant for the control of receptor signalling responses (3). Agonist-induced GLP-1R phosphorylation is mediated by GPCR kinases (GRKs) (4) and occurs predominantly on serine residues within the receptor’s intracellular C-terminal tail, with three serine (Ser) doublets previously identified as GLP-1R phospho-sites in positions 441/442, 444/445, and 451/452 (5) of the receptor. These receptor phosphorylation events are thought to promote β-arrestin recruitment, receptor desensitisation, and internalisation, thereby shaping both the amplitude and the duration of GLP-1R signalling. However, most existing data on the effect of GLP-1R phosphorylation in regulating receptor signalling responses are indirect (4, 6, 7), and/or have been collected from heterologous expression systems (4, 5, 7, 8), so that the validity of these observations in physiologically relevant models with endogenous expression of the receptor and its downstream signalling mediators [GRKs, β-arrestins, phosphodiesterases (PDEs), endosomal trafficking regulators, etc.] is currently unknown. Emerging evidence indicates that distinct phosphorylation patterns, or “barcodes”, can influence GPCR engagement with β-arrestins and determine signalling bias (9), potentially contributing to the disparity of pharmacological and therapeutic profiles observed among different GLP-1R agonists (10). This makes the modelling of the mechanisms and functional consequences of GLP-1R phosphorylation in relevant cell systems essential for the meaningful elucidation of receptor signalling dynamics to guide the rational design of next-generation incretin-based therapeutics.

Here, we sought to address this knowledge gap by modelling the trafficking and signalling behaviours of GLP-1R phospho-site null mutants in pancreatic β-cells, uncovering the C-terminal Ser451/452 doublet as critical for the recruitment of GRK2 to the receptor, and revealing an unexpected positive correlation between GLP-1R phosphorylation and β-cell cAMP production following receptor activation by its natural ligand, GLP-1.

## Materials and Methods

### Peptides and drugs

GLP-1 (7-36) acetate (HY-P0054) were purchased from MedChemExpress. Rolipram (557330), forskolin (F6886), and 3-isobutyl-1-methylxanthine (IBMX) (I5879) were purchased from Sigma-Aldrich.

### Cell culture

INS-1 832/3 cells with endogenous rat *Glp1r* deleted by CRISPR/Cas9 (11) (referred to as INS-1 832/3 GLP-1R KO cells, kind gift from Dr Jacqueline Naylor, Astra Zeneca) were derived from parental male rat insulinoma INS-1 832/3 cells (Prof Christopher Newgard, Duke University, USA). Cells were cultured in RPMI-1640 media with 11 mM D-glucose (Gibco) supplemented with 10% fetal bovine serum (FBS), 10 mM HEPES, 1 mM sodium pyruvate, 50 μM β-mercaptoethanol, and 1% penicillin/streptomycin at 37°C in a 95% O_2_ / 5% CO_2_ incubator.

### Human phospho-site mutant GLP-1R construct generation

SNAP/FLAG-tagged human GLP-1R wild-type (WT) plasmid (Cisbio), subsequently referred to as SNAP/FLAG-hGLP-1R, was altered by site-directed mutagenesis with the PfuUltra II Fusion High-fidelity DNA Polymerase (Agilent) as per the manufacturer’s protocol to modify serine to alanine doublets at positions 441/442, 444/445, and 451/452 of the receptor to generate individual serine doublet mutants (referred to as S441/2A, S444/5A and S451/2A). To generate the triple AA mutant with all three serine doublets mutated to alanine, the S441/2A mutant was used as a template to introduce serial alanine modifications at positions 451/452 and 444/445 using mutagenesis primers specifically designed to preserve the previous serine to alanine mutations. SNAP/FLAG-hGLP-1R-NanoLuc was previously generated in house by polymerase chain reaction (PCR) cloning of the NanoLuciferase sequence from pcDNA3.1-ccdB-NanoLuc (kind gift from Prof Mikko Taipale; Addgene plasmid #87067) onto the C-terminal end of SNAP/FLAG-hGLP-1R followed by site-directed mutagenesis of the GLP-1R stop codon. Individual serine doublet and triple AA phospho-site alanine mutations were introduced at the SNAP/FLAG-hGLP-1R-NanoLuc plasmid as described above for the SNAP/FLAG-hGLP-1R construct. Mutagenesis of selected serine to alanine residues was confirmed by Sanger sequencing (Genewiz). Mutagenesis and sequencing primer sequences are listed in Table 1.

### SNAP/FLAG-hGLP-1R serine phosphorylation analysis by immunoprecipitation

INS-1 832/3 GLP-1R KO cells were seeded in 6-well plates 24 hours before transfection with 2 µg WT or phospho-site mutant SNAP/FLAG-hGLP-1R constructs using Lipofectamine 2000 (Thermo Fisher Scientific) as per the manufacturer’s instructions, with all subsequent transfections in the study also performed with this reagent. Transfected cells were seeded onto vehicle and GLP-1 treatment wells 24 hours post-transfection and treated 24 hours post-seeding with vehicle or 100 nM GLP-1 for 5 minutes in complete medium. Following treatment, cells were washed 1x in PBS before being lysed in ice-cold lysis buffer (50 mM Tris-HCl, pH 7.4, 150 mM NaCl supplemented with 1 mM EDTA, 1% Triton X-100, protease and phosphatase inhibitor cocktails), with lysates gently rocked for 15 minutes at 4°C before centrifugation at 13,000 g for 5 minutes at 4°C. Supernatants were recovered and incubated for 2 hours under gentle rotation at room temperature (RT) with Anti-FLAG M2 Magnetic beads (Merck) to immunoprecipitate SNAP/FLAG-hGLP-1R according to the manufacturer’s protocol.

Following immunoprecipitation, beads were resuspended in 2x urea loading buffer (20 mM Tris-HCl, pH 6.8, 5% SDS, 8 M urea, 100 mM dithiothreitol, and 0.02% bromophenol blue) at 1:1 volume/volume ratios and incubated at 37°C for 10 minutes to elute the immunoprecipitated proteins from the beads. Beads were separated on a magnetic rack, and supernatants resolved by SDS-PAGE (Bio-Rad). Proteins were transferred to PVDF membranes (Immobilon-P, 0.45-μm pore size, Sigma-Aldrich) using a wet transfer Western blot system (Bio-Rad) and membranes blocked in 5% BSA in 1x TBS/Tween (20 mM Tris-HCl, 150 mM NaCl, 0.1% Tween-20) before incubation with a mouse monoclonal anti-pan-phospho-serine primary antibody (M380B, Biolegend, 1:1,000) also in 5% BSA in 1x TBS/Tween followed by 3x washes in 1x TBS/Tween and incubation with goat anti-mouse HRP secondary antibody (ab205719, Abcam, 1:5,000) before being developed using the Clarity Western enhanced chemiluminescence substrate system (Bio-Rad) in a ChemiDoc imaging system (Bio-Rad). Membranes were subsequently stripped and incubated with anti-SNAP tag rabbit polyclonal antibody (P9310S, New England Biolabs, 1:1,000) followed by goat anti-rabbit HRP secondary antibody (ab6721, Abcam, 1:2,000) to detect total level of immunoprecipitated SNAP/FLAG-hGLP-1R per sample. Specific band densities were quantified using Fiji ImageJ and phospho-serine bands normalised to SNAP levels per lane.

### GLP-1R surface expression and internalisation assays by high-content microscopy

The assays were performed as previously described (12). INS-1 832/3 GLP-1R KO cells were reverse-transfected onto poly-L-lysine-coated, Optical-Bottom black 96-well plates with 0.2 µg WT or serine doublet mutant SNAP/FLAG-hGLP-1R constructs. The next day, a 10-minute labelling was performed with the SNAP-Surface cleavable probe BG-S-S-649 (kind gift from Dr Ivan Corrêa Jr., New England Biolabs) in complete medium at 37°C. After washing in PBS, cells were treated with vehicle or 100 nM GLP-1 for 10 minutes at 37°C in complete medium before being washed 1x in cold PBS, with all subsequent steps performed on ice. Cells were then treated with 100 mM reducing agent sodium 2-mercaptoethanesulfonate (Mesna) in TNE buffer (20 mM Tris, 150 mM NaCl, 1 mM EDTA, pH 8.6) or with TNE buffer alone for 5 minutes before being washed 1x in PBS and imaged in Hanks’ Balanced Salt Solution (HBSS) buffer on a Nikon Ti2E wide-field microscope with a light-emitting diode (LED) light source and a 20x/0.75 NA objective, with several phase contrast and epifluorescence images acquired per well using corresponding filters. Internalised receptor from GLP-1-treated cells was quantified from cell-containing regions as determined from the phase contrast images using PHANTAST (13) and used to determine receptor internalisation as previously described (12). Internalisation was expressed as a percentage of total background-subtracted surface labelled receptor signal from non-Mesna-treated vehicle wells, a value that was also used to estimate cell surface expression levels of the SNAP-tagged receptors.

### Plasma membrane diffusion assay by Raster imaging correlation spectroscopy (RICS)

INS-1 832/3 GLP-1R KO cells were transfected with 0.5 µg WT or triple AA phospho-null mutant SNAP/FLAG-hGLP-1R constructs before seeding onto 35-mm glass-bottom dishes (MatTek Corporation) and labelling with membrane-impermeable SNAP-Surface Alexa Fluor 647 probe (S9136S, New England Biolabs) for 12 minutes at 37°C in complete media. Cells were subsequently washed and imaged in RMPI-1640 media without phenol red (Gibco) using a 100x/1.4 NA oil objective on a Leica Stellaris 8 STED FALCON microscope from the Imperial Facility for Ligh Microscopy (FILM), using previously described confocal settings (14). Image analysis was performed as previously described (15, 16). In brief, cells were imaged at the plasma membrane under vehicle conditions as well as following stimulation with 100 nM GLP-1 with an image size of 256×256 pixels and 80 nm pixel size for 200 consecutive frames. RICS analysis was performed to determine the diffusion coefficient of receptors in vehicle *versus* GLP-1-stimulated conditions using the SimFCS 4 Software (Global Software, G-SOFT Inc.). Three different regions of interest of 32×32 pixels per image and a moving average of 10 was applied to avoid any artefacts due to cellular motion or very slow-moving particles. Average intensity, intensity plots, 2D autocorrelation maps and 3D autocorrelation fits were generated per condition.

### cAMP generation and β-arrestin 2 recruitment assays

INS-1 832/3 GLP-1R KO cells were transfected with 0.5 µg WT or phospho-site mutant SNAP/FLAG-hGLP-1R constructs before being seeded onto 35-mm glass-bottom dishes (MatTek Corporation) and transduced with the Green Up cADDis cAMP (Big Sky) biosensor for cAMP assays, or the Green Fluorescence Borealis Arrestin biosensor in a BacMam vector (Montana Molecular) as per the manufacturer’s instructions. 12 hours post-transduction, cells were labelled with SNAP-Surface Alexa Fluor 647, washed 1x in PBS and imaged in RPMI-1640 media without phenol red using a Nikon Eclipse Ti microscope with an ORCA-Flash 4.0 camera (Hamamatsu) and Metamorph software (Molecular Devices) with a 20x air objective and 488 and 647 nm lasers. A single image was initially captured at both λ_ex_ 488 and 647 nm to assess SNAP-tagged receptor expression and biosensor transduction distributions, followed by images acquired every 6 seconds at λ_ex_ 488 nm to assess cAMP or β-arrestin 2 responses on a 37°C stage. A baseline reading was recorded for 1 minute prior to stimulation with 100 nM GLP-1 for 5 minutes, and 10 μM forskolin + 100 μM IBMX added for the last 2 minutes of the acquisition to record maximal responses. Raw fluorescent intensities were extracted in Fiji ImageJ to plot average intensity traces and normalised to average baseline responses prior to calculation of AUCs for the GLP-1 stimulation period in GraphPad Prism 10.2.1. Bias between cAMP generation and β-arrestin 2 recruitment was calculated using the ratios of the mean AUCs for each parameter with errors propagated according to the formula 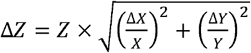, with 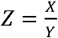 and x and y representing the respective mean AUC values.

### GRK2-Venus recruitment assays

INS-1 832/3 GLP-1R KO cells seeded onto 12-well plates were co-transfected with 0.5 µg each of GRK2-Venus (17) (kind gift from Prof Meritxell Canals, University of Nottingham) and either WT or phospho-site mutant SNAP/FLAG-hGLP-1R-NanoLuc constructs. 24 hours post-transfection, cells were detached, resuspended in NanoGlo Live Cell Reagent (Promega) with furimazine (1:20 dilution) and seeded onto 96-well opaque white half-area plates. Baseline luminescent signals were recorded every minute at 460 nm (NanoLuc emission peak) and 535 nm (Venus emission peak) for 5 minutes at 37°C, followed by 30 minutes with or without addition of 100 nM GLP-1. Raw relative light unit (RLU) readings were expressed as the ratio of Venus to Nanoluc signals, and baseline-corrected BRET ratios normalised to vehicle readings to determine the agonist-induced effect. AUCs from response curves were calculated using Graphpad Prism 10.2.1.

### Statistical analyses

All statistical analyses were performed using GraphPad Prism 10.2.1. Data are expressed as mean ± SEM. Statistical significance was calculated using two-tailed Student’s t-test, paired or unpaired depending on the manner of acquisition, for experiments with two variables, or by one-way analysis of variance (ANOVA) for experiments with three or more variables, with comparison to a single condition or across conditions depending on the nature of parameters measured, with either Dunnett’s or Šidák’s post-hoc tests as indicated in the figure legends. Outlier detection was performed using the ROUT method with a Q value of 10%.

## Results

As mentioned above, the human GLP-1R phospho-sites have been previously mapped to three serine doublets located in the C-terminal tail of the receptor (5) (Fig. 1A). Based on this, we have employed a site-directed mutagenesis approach to generate a phospho-site mutant construct for each serine doublet in the human receptor, by mutating each doublet to alanine (mutants referred to as S441/2A, S444/5A and S451/2A). SNAP/FLAG-tagged wildtype (WT) or serine doublet mutant human GLP-1R constructs were expressed in INS-1 832/3 rat insulinoma cells with the endogenous rat receptor previously knocked out by CRISPR/Cas9 (11) (INS-1 832/3 GLP-1R KO cells), and WT or mutant receptors subsequently purified from these cells by immunoprecipitation under vehicle conditions or following a 5-minute stimulation with 100 nM GLP-1. Western blot detection of serine phosphorylation levels in the immunoprecipitated receptor fraction using an anti-pan-phospho-serine antibody confirmed that the WT GLP-1R is indeed serine phosphorylated in response to GLP-1 exposure (Fig. 1B, C). GLP-1-induced receptor serine phosphorylation was unexpectedly completely blunted, rather than partially reduced, in all three individual serine doublet mutants compared to the WT GLP-1R. We next performed combined site-directed mutagenesis to generate a receptor construct with all three serine doublets mutated to alanine (triple AA phospho-null mutant). As expected, following the same immunoprecipitation approach as above, we measured a complete loss of serine phosphorylation in response to GLP-1 stimulation in the triple AA mutant compared to the WT receptor (Fig. 1D, E).

**Figure 1.**
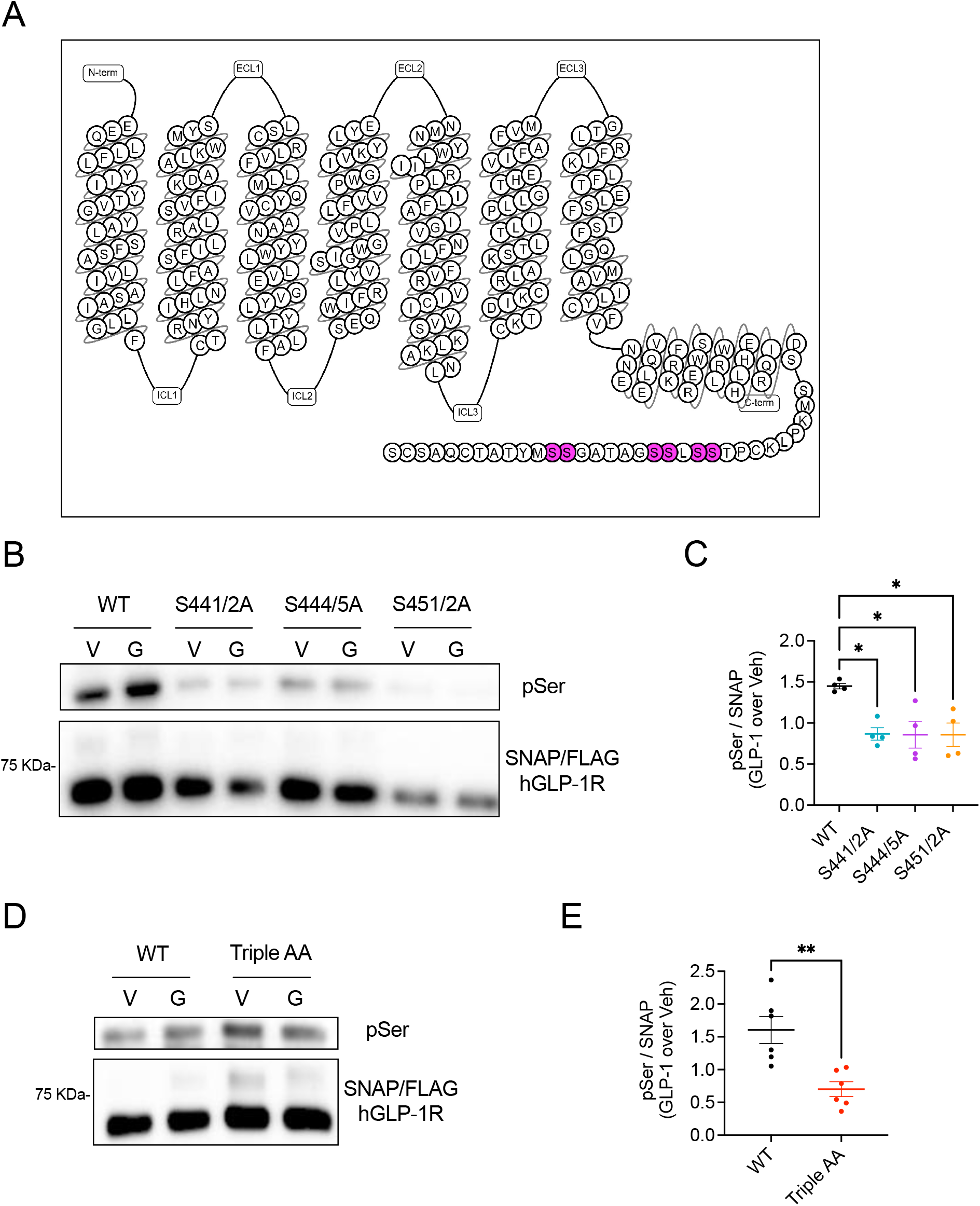
GLP-1-dependent human GLP-1R phosphorylation in β-cells. (A), Snake plot of human GLP-1R primary sequence depicting C-terminal tail serine doublets at positions 441/442, 444/445, and 451/452, as highlighted in magenta; adapted from GPCRdb. (B) and (C), Western blot analysis of serine phosphorylation levels in WT *versus* individual serine doublet mutant SNAP/FLAG-hGLP-1Rs immunoprecipitated from INS-1 832/3 GLP-1R KO cells after vehicle (V) or 5-minute stimulation with 100 nM GLP-1 (G); Representative blots (B) and quantification of phospho-serine (pSer) over SNAP levels normalised to vehicle (Veh) (C); *n*=4 biologically independent experiments. (D) and (E), As in (B) and (C) for WT *versus* triple AA phospho-null SNAP/FLAG-hGLP-1R; *n*=6 biologically independent experiments. Data are mean ± SEM; *p<0.05, **p<0.01, by paired t-test or one-way ANOVA with Dunnett’s post-hoc test.

Having assessed the relevance of the three C-terminal serine doublets on the phosphorylation of the receptor in response to agonist stimulation, we next investigated the effect of mutating each doublet to alanine on the capacity of the receptor to traffic to the plasma membrane and therefore be available for agonist binding, as well as its propensity to undergo agonist-induced endocytosis in INS-1 832/3 GLP-1R KO cells. All three individual serine doublet mutant receptors exhibited similar or even increased cell surface levels compared to WT GLP-1R, indicating that these serine to alanine substitutions in the C-terminal tail do not hinder the receptor biosynthesis or its subsequent targeting to the plasma membrane (Fig. 2A). While we did not observe any significant changes, there was a general tendency for all three serine doublet mutants to display reduced internalisation in response to GLP-1 stimulation compared to the WT GLP-1R (Fig. 2B). We next analysed these parameters on the triple AA phospho-null mutant receptor and again found similar, non-significantly raised cell surface receptor levels for the mutant compared to the WT GLP-1R (Fig. 2C). This time, however, the triple AA phospho-null mutant exhibited significantly reduced internalisation propensity in response to GLP-1 stimulation *versus* the WT receptor (Fig. 2D).

**Figure 2.**
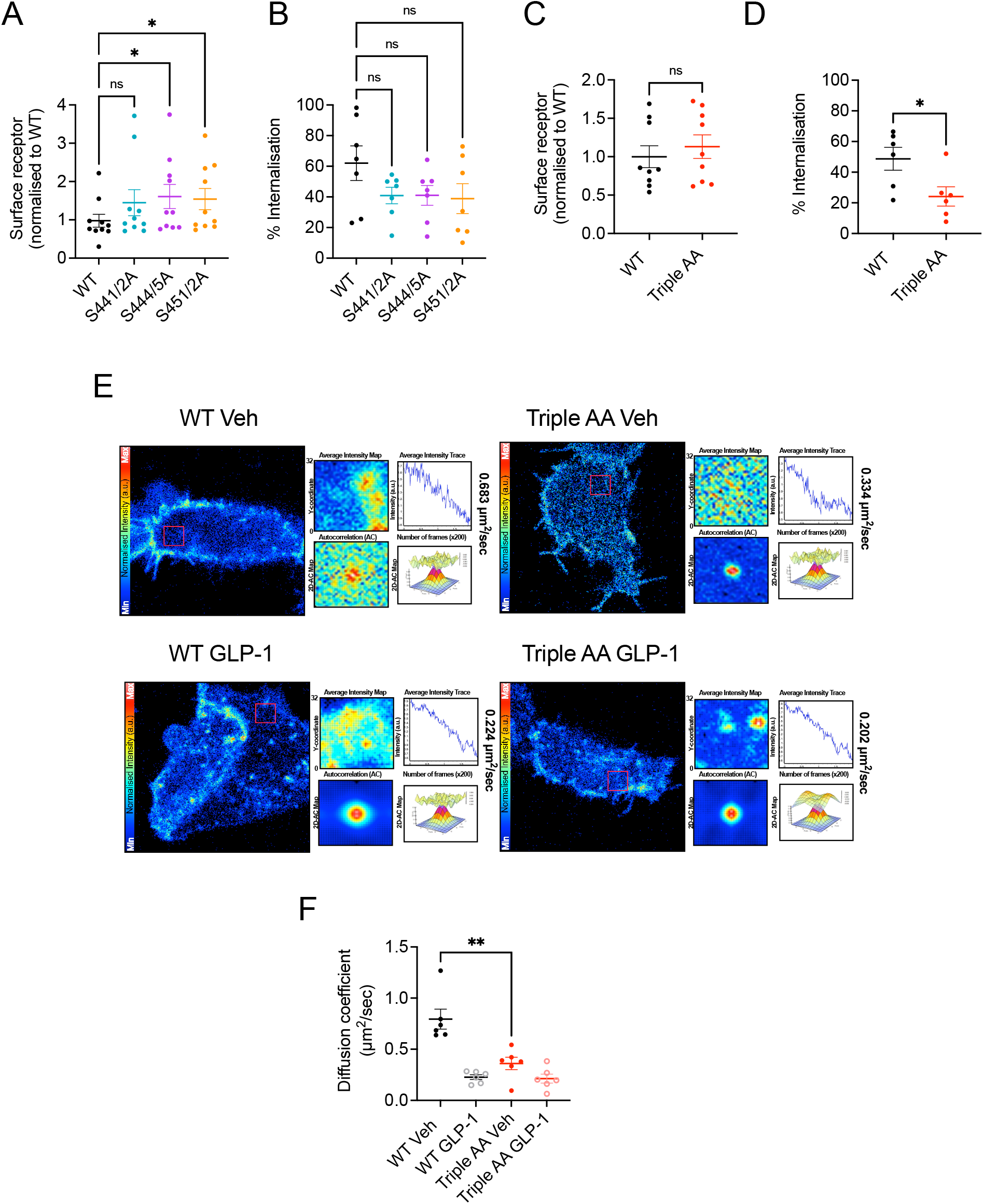
Surface expression levels and GLP-1-induced internalisation of WT *versus* phospho-mutant human GLP-1R in β-cells. (A), WT *versus* individual serine doublet mutant SNAP/FLAG-hGLP-1R surface expression levels, quantified by high-content microscopy of SNAP-Surface 647-labelled INS-1 832/3 GLP-1R KO cells; data normalised to WT receptor levels; *n*=10 biologically independent experiments. (B), Percentage of GLP-1-induced GLP-1R internalisation, quantified by high-content microscopy (Mesna) assay, in WT *versus* individual serine doublet mutant SNAP/FLAG-hGLP-1Rs expressed in INS-1 832/3 GLP-1R KO cells; *n*=7 biologically independent experiments. (C), As in (A) for WT *versus* triple AA phospho-null mutant SNAP/FLAG-hGLP-1R; *n*=9 biologically independent experiments. (D), As in (B) for WT *versus* triple AA phospho-null mutant SNAP/FLAG-hGLP-1R; *n*=6 biologically independent experiments. (E) and (F), Representative images (E) and quantification of plasma membrane diffusion rates (F) obtained by RICS analysis of SNAP-Surface 647-labelled WT *versus* triple AA phospho-null mutant SNAP/FLAG-hGLP-1Rs expressed in INS-1 832/3 GLP-1R KO cells treated with vehicle (Veh) *versus* 100 nM GLP-1; diffusion coefficients for each representative ROI indicated per image, with corresponding intensity traces, 2D and fitted 3D autocorrelation maps included; *n*=6 biologically independent experiments. Data are mean ± SEM; ns, non-significant, *p<0.05, **p<0.01, by paired t-test or one-way ANOVA with Šidák’s post-hoc test.

We next investigated if the triple AA phospho-null mutant receptor would display any differences in its dynamic behaviour at the plasma membrane by analysing WT *versus* mutant receptor membrane diffusion rates by Raster image correlation spectroscopy (RICS), a technique that allows the assessment of molecular dynamics from fluorescence confocal images (18), which we have used in the past to assess the dynamic behaviour of incretin receptors (16, 19). RICS analysis of WT receptors in INS-1 832/3 GLP-1R KO cells showed, as previously observed (19), that the GLP-1R is highly dynamic under vehicle conditions but undergoes sharply reduced diffusion, typically associated with increased receptor clustering, following GLP-1 stimulation (Fig. 2E, F). In contrast, the triple AA phospho-null mutant GLP-1R exhibited highly reduced plasma membrane diffusion at vehicle conditions *versus* the WT receptor, with a similarly low diffusion rate in both following GLP-1 exposure.

To follow up these investigations, we next examined the capacity of the individual serine doublet mutant receptors to elicit downstream signalling responses in INS-1 832/3 GLP-1R KO cells, including analyses of GLP-1-induced cAMP generation and β-arrestin 2 recruitment to the receptor. Individual serine mutant GLP-1Rs displayed a tendency towards reduced GLP-1-induced cAMP generation without reaching statistical significance when compared to the WT receptor (Fig. 3A, B). No differences were found between WT and individual serine doublet mutants on the capacity to recruit β-arrestin 2 downstream of GLP-1 stimulation (Fig. 3C, D).

**Figure 3.**
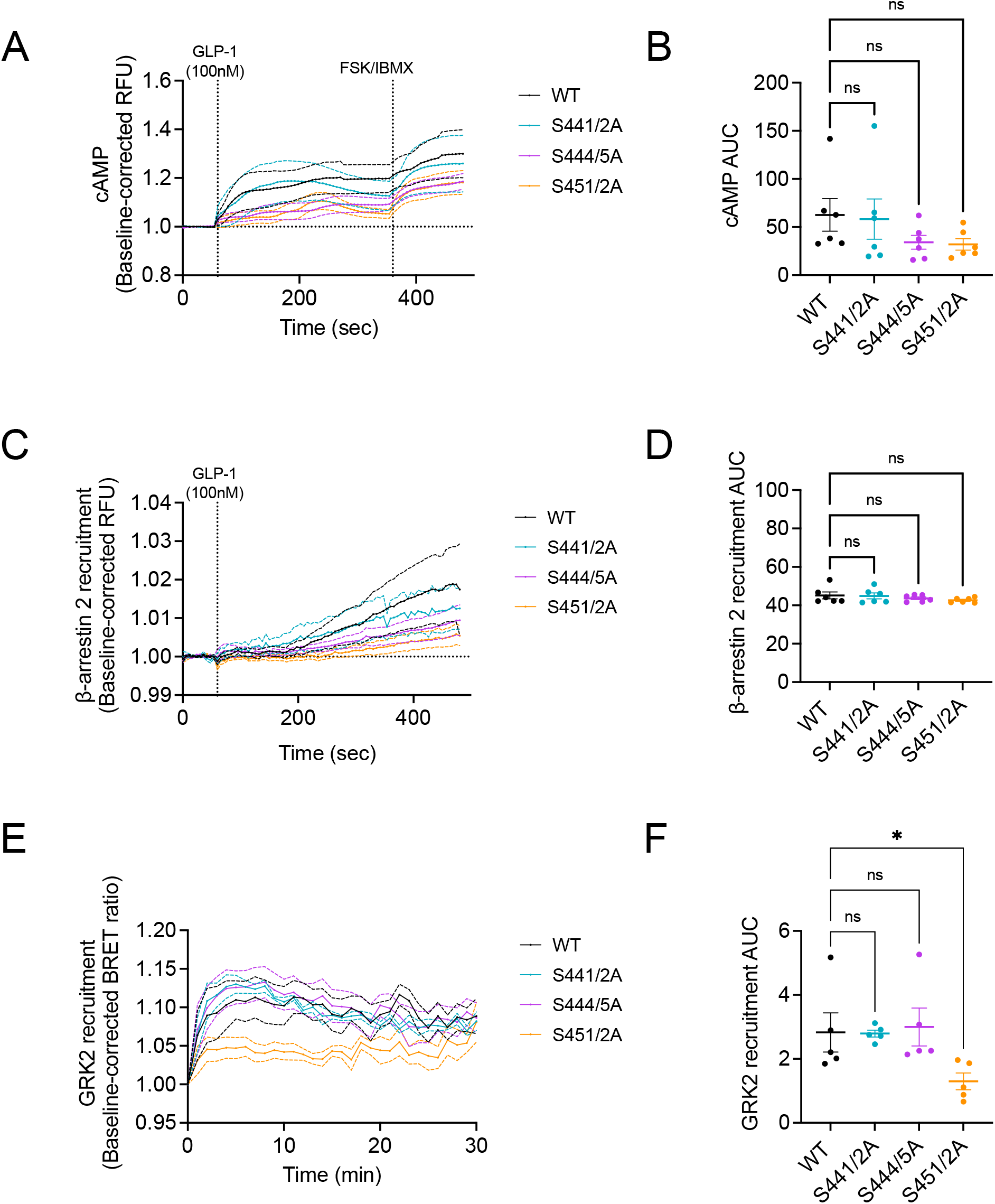
Signalling responses of WT *versus* individual serine doublet mutant human GLP-1Rs in β-cells. (A) and (B), cAMP kinetic responses (A) and corresponding AUC (B) of WT *versus* individual serine doublet mutant SNAP/FLAG-hGLP-1Rs expressed in INS-1 832/3 GLP-1R KO cells in response to 100 nM GLP-1 stimulation; assessed by spinning disk microscopy with the fluorescent cAMP biosensor cADDis; responses normalised to baseline, maximal responses with 10 μM forskolin (FSK) + 100 μM IBMX assessed at the acquisition end, FSK + IBMX period not included in AUC quantification; *n*=6 biologically independent experiments. (C) and (D), β-arrestin 2 recruitment kinetic responses (C) and corresponding AUC (D) for WT *versus* individual serine doublet mutant SNAP/FLAG-hGLP-1Rs in INS-1 832/3 GLP-1R KO cells in response to 100 nM GLP-1 stimulation; assessed by spinning disk microscopy with the fluorescent β-arrestin 2 recruitment biosensor Borealis; responses normalised to baseline; *n*=6 biologically independent experiments. (E) and (F), GRK2-Venus recruitment kinetic responses (E) and corresponding AUC (F) for WT *versus* individual serine doublet mutant SNAP/FLAG-hGLP-1R-Nanoluc constructs in INS-1 832/3 GLP-1R KO cells in response to 100 nM GLP-1 stimulation; data obtained by NanoBRET assay; baseline-corrected results normalised to vehicle; *n*=5 biologically independent experiments. Data are mean ± SEM; ns, non-significant, *p<0.05 by one-way ANOVA with Dunnett’s post-hoc test.

Finally, we also examined the capacity of each mutant to recruit GRK2, a GPCR kinase known to be recruited to active GLP-1R and to regulate GLP-1R-mediated secretory responses in β-cells (20). These experiments confirmed that GRK2 is indeed recruited to the WT receptor downstream of GLP-1 stimulation and identified the third serine doublet at position 451/452, but not the other two doublets, as required for the recruitment of GRK2 to GLP-1-stimulated GLP-1R in our β-cell line (Fig. 3E, F).

Having analysed the functional impact of each serine doublet mutant receptor individually, we next examined the signalling profile of the triple AA phospho-null mutant in INS-1 832/3 GLP-1R KO cells. Results showed a clear defect in cAMP generation downstream of GLP-1 stimulation for the triple AA mutant compared to the WT receptor in these cells (Fig. 4A, B). Additionally, while we measured a near significant decrease in GLP-1-induced β-arrestin 2 recruitment to the mutant compared to the WT GLP-1R (Fig. 4C, D), this subtle effect was a lot less pronounced than the previously observed loss of cAMP generation, so that a bias calculation between both parameters indicated that, at least in our cells, the triple AA phospho-null mutant exhibits biased signalling towards β-arrestin 2 recruitment *versus* cAMP generation compared to WT GLP-1R (Fig. 4E).

**Figure 4.**
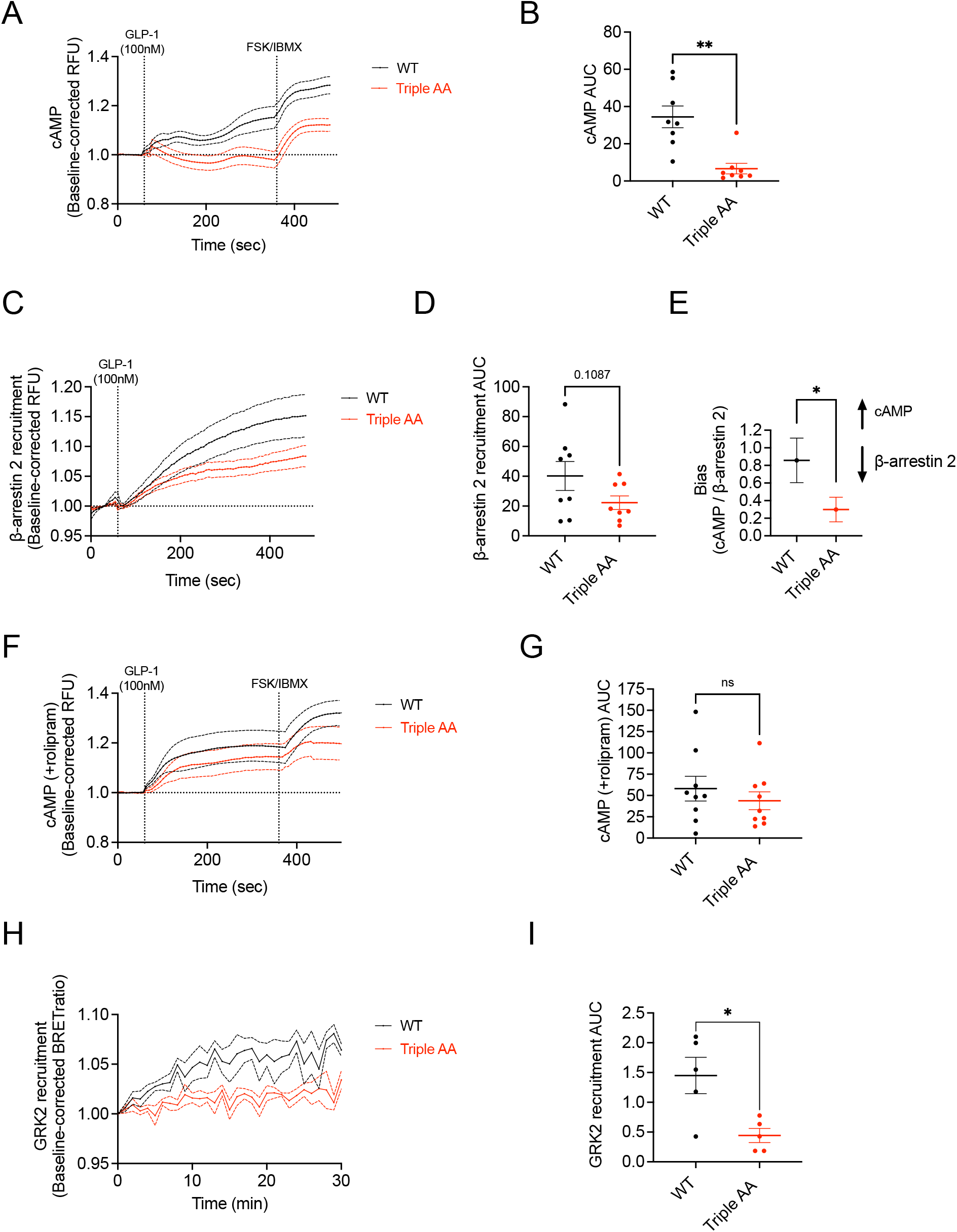
Signalling responses of WT *versus* triple AA phospho-null mutant human GLP-1R in β-cells. (A) and (B), cAMP kinetic responses (A) and corresponding AUC (B) of WT *versus* triple AA phospho-null mutant SNAP/FLAG-hGLP-1R expressed in INS-1 832/3 GLP-1R KO cells in response to 100 nM GLP-1 stimulation; assessed by spinning disk microscopy with the fluorescent cAMP biosensor cADDis; responses normalised to baseline, maximal responses with 10 μM forskolin (FSK) + 100 μM IBMX assessed at the acquisition end, FSK + IBMX period not included in AUC quantification; *n*=8 biologically independent experiments. (C) and (D), β-arrestin 2 recruitment kinetic responses (C) and corresponding AUC (D) for WT *versus* triple AA phospho-null mutant SNAP/FLAG-hGLP-1R in INS-1 832/3 GLP-1R KO cells in response to 100 nM GLP-1 stimulation; assessed by spinning disk microscopy with the fluorescent β-arrestin 2 recruitment biosensor Borealis; responses normalised to baseline; *n*=8 biologically independent experiments. (E), Bias between cAMP generation and β-arrestin 2 recruitment in response to 100 nM GLP-1 stimulation for WT *versus* triple AA phospho-null mutant SNAP/FLAG-hGLP-1R expressed in INS-1 832/3 GLP-1R KO cells, calculated from data in (B) and (D). (F) and (G), As in (A) and (B) in the presence of 10 µM PDE4 inhibitor rolipram; *n*=9 biologically independent experiments. (H) and (I), GRK2-Venus recruitment kinetic responses (H) and corresponding AUC (I) for WT *versus* triple AA phospho-null mutant SNAP/FLAG-hGLP-1R-Nanoluc in INS-1 832/3 GLP-1R KO cells in response to 100 nM GLP-1 stimulation; data obtained by NanoBRET assay; baseline-corrected results normalised to vehicle; *n*=5 biologically independent experiments. Data are mean ± SEM; ns, non-significant, *p<0.05 by paired t-test, or by p value calculation from t score of mean AUC ratio and error propagation in (E).

To gain a deeper understanding of the mechanism involved in the loss of cAMP generation observed in the triple AA phospho-null mutant receptor, we repeated the cAMP assays in INS-1 832/3 GLP-1R KO cells expressing WT or triple AA mutant receptors pre-treated with the PDE4-specific inhibitor rolipram (21). We have in the past shown that PDE4 inhibition can restore normal levels of GLP-1R-dependent cAMP generation in islets with increased receptor desensitisation due to loss of β-arrestin 2 expression (22). Here again, rolipram treatment was able to restore, at least partially, the cAMP response of the triple AA phospho-null mutant so that this was no longer significantly reduced when compared to the WT receptor (Fig. F, G).

Finally, we also assessed the capacity of the triple AA phospho-null mutant receptor to recruit GRK2 downstream of GLP-1 stimulation and we again observed a significant loss of in the recruitment of this kinase in the mutant *versus* the WT receptor (Fig. 4H, I), as previously seen in Fig. 3E, F for the S451/2A doublet mutant receptor.

## Discussion

GPCR phosphorylation is a crucial PTM involved in the control of cell signalling by regulating receptor desensitisation and internalisation propensities, recruitment of β-arrestins, and redirecting receptors towards specific signalling pathways (23). Previous investigations focusing on the control of GLP-1R signalling responses by this PTM have however yielded contradictory reports. An early study in fibroblasts, which identified the three serine doublets in the C-terminal tail of the receptor investigated here as relevant phospho-sites downstream of GRK action, showed a positive correlation between receptor phosphorylation and homologous desensitisation and linked receptor phosphorylation state with its propensity for internalisation (5). Conversely, a recent study in HEK cells did not show any significant effect in cAMP accumulation and only minimal effect on internalisation of phospho-site mutant GLP-1Rs (8). Both studies implied reductions in the recruitment of β-arrestins to the phospho-site mutant receptors, which, confusingly, were only mildly altered in mutant GLP-1Rs where the whole C-terminal tail is substituted by that of the low β-arrestin-recruiting GIP receptor (GIPR) (24). All these studies were however performed in non-β-cell lines, which could carry serious caveats related to cell-type specific patterns of expression of receptors, G proteins, GRKs, and β-arrestin isoforms (25), hindering the interpretation of effects related to changes in the engagement of these intertwined regulatory factors.

Our investigation employs the rat INS-1 832/3 β-cell line, which naturally expresses the GLP-1R (26) and responds to GLP-1 by potentiating glucose-induced insulin secretion (GSIS) and increasing intracellular cAMP levels (19), making it a valuable model for studying the effect of GLP-1R phospho-site mutants on β-cell signalling outputs. By expressing the human WT *versus* individual serine doublet or triple AA mutant receptors on a rat GLP-1R null background, we sought to understand the impact of GLP-1R phosphorylation on the control of its signalling on a β-cell context. Reassuringly, we could demonstrate a ~1.5-fold increase in serine phosphorylation of the WT receptor by acute (5-minute) stimulation with GLP-1. Interestingly however, the receptor was not completely unphosphorylated under vehicle conditions, suggesting that there is some basal level of GRK recruitment and phosphorylation present under unstimulated conditions, with the receptor potentially cycling between phosphorylated and unphosphorylated states by a currently unknown mechanism. Also surprisingly, the GLP-1 potentiation of receptor serine phosphorylation was completely lost rather than partially reduced for all three individual serine doublet mutants compared to the WT GLP-1R, indicating that the receptor requires the presence of all three C-terminal serine doublets for sustained phosphorylation downstream of GLP-1 stimulation, at least as far as our method of detection of serine-phosphorylated receptor allows us to measure. This observation suggests that each serine doublet might play a specialised function in the phosphorylation process, for example by recruiting specific GRKs, as shown in the present study for GRK2, which requires the presence of the third serine doublet at position 451/452 of the C-terminal tail to be recruited to the receptor downstream of GLP-1 stimulation.

A further observation of this study relates to the capacity of the receptor serine doublet phospho-sites to regulate its dynamic behaviour under non-stimulated conditions, as observed in our RICS analysis of receptor plasma membrane diffusion, which demonstrated significantly reduced basal diffusion rate of the triple AA phospho-null mutant compared to the WT receptor under vehicle conditions. The reason for this difference is currently unclear but could potentially be related to changes in the capacity of the phospho-null mutant GLP-1R C-terminal tail to establish interactions with specific lipids and/or modulate receptor oligomerisation states, two processes where the receptor tail has previously been implicated (27). Regardless of the mechanism leading to reduced diffusion of the unstimulated phospho-null receptor, this phenomenon, together with the previous observation of some degree of phosphorylation at the WT receptor under vehicle conditions, indicates the existence of a much more dynamic cycling between phosphorylated and unphosphorylated receptor states, present already in the apo-state, which could potentially involve interactions not only with specific GRKs but also with phosphatases that could contribute to the regulation of the overall degree of receptor phosphorylation, explaining why the C-terminal serine phospho-sites have the capacity to influence the behaviour of the receptor even in the absence of agonist stimulation.

A further observation from the present study is the link between receptor serine phosphorylation and optimal receptor endocytosis, as previously shown in (5), although this effect appears to be only partial in our β-cell system. This observation was assumed in the past to be linked to the recruitment of β-arrestins to phosphorylated receptors given their role in facilitating receptor internalisation, but this hypothesis must be reconciled with the current notion of lack of β-arrestin requirement for efficient GLP-1R endocytosis (4, 22, 27). Agreeably, we could only detect a non-significant tendency for reduced β-arrestin 2 recruitment in our triple AA phospho-null mutant receptor despite significantly reduced internalisation in response to GLP-1 stimulation, with this effect more likely ascribed to the observed reduction in GRK2 recruitment, in agreement with a previous report highlighting GRK expression as required for GLP-1R internalisation (4). Our observation of only a tendency for reduced β-arrestin 2 recruitment at the phospho-null mutant receptor is also in agreement with a previous report highlight the biphasic mode of β-arrestin 2 – GLP-1R interaction, with a phosphorylation-independent as well as a phosphorylation-dependent component (28).

By far the most unexpected result of the present study is the defect in cAMP generation that we have measured downstream of GLP-1 stimulation for the triple AA phospho-null mutant compared to the WT receptor in our β-cell system. This effect, already hinted at in the individual serine doublet mutants, contrasts with the increase in cAMP associated with reduced homologous desensitisation of the phospho-null mutant previously observed in a heterologous system (5). While further investigations are required to fully elucidate why the phospho-null mutant GLP-1R is unable to sustain GLP-1-induced cAMP signalling in β-cells, alterations in the recruitment of specific PDEs, which can potentially be cell-type specific and therefore not always replicated in non-β-cell systems, are likely to play an important role, as shown here by the capacity of the PDE4-specific inhibitor rolipram to restore cAMP responses from the triple AA phospho-null mutant receptor.

In conclusion, this study has analysed the effect of mutating three serine doublets in the C-terminal tail of the GLP-1R to alanine, both individually and combined, in a β-cell model (see Fig. 5 for a graphical summary of the main results of the study), confirming their requirement for agonist-induced GLP-1R phosphorylation, and identifying the third serine doublet at position 451/452 as required for the recruitment of GRK2 to the stimulated receptor, as well as unveiling a previously unknown positive correlation between receptor phosphorylation and its capacity for cAMP generation downstream of GLP-1 stimulation in β-cells.

**Figure 5.**
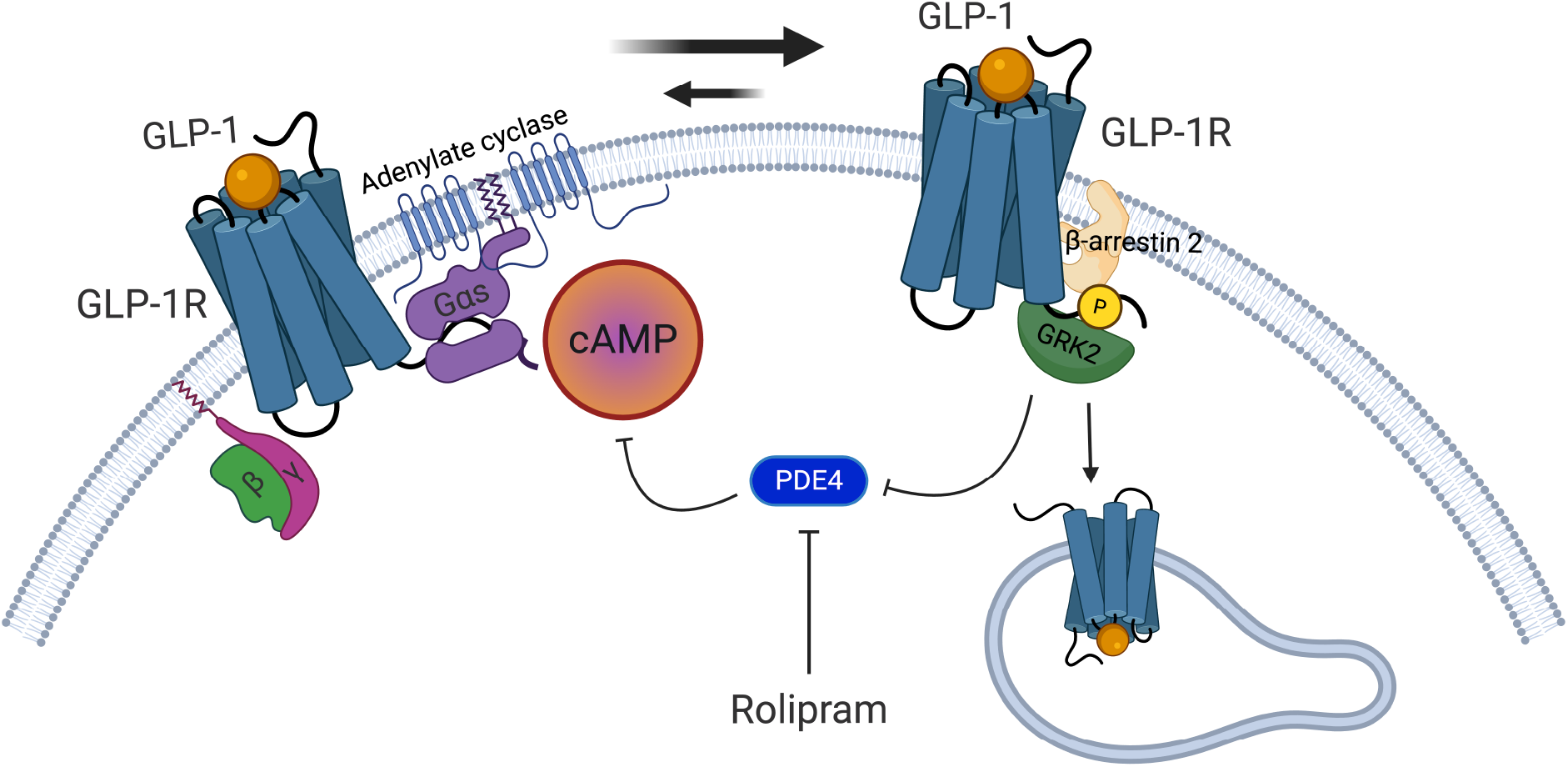
Graphical summary of impact of human GLP-1R serine phosphorylation on receptor trafficking and signalling responses in β-cells. Following GLP-1 stimulation, the GLP-1R C-terminal tail is phosphorylated in three serine doublets at the receptor C-terminal tail, with all three doublets required for GLP-1-induced GLP-1R serine phosphorylation but only Ser451/452 doublet required for GRK2 recruitment to the receptor. GLP-1R triple AA phospho-null mutant displays reduced plasma membrane diffusion rates under basal conditions, suggesting intrinsic effects of phospho-site mutations prior to receptor activation. Other changes include reduced GLP-1-induced receptor internalisation and a tendency for reduced β-arrestin 2 recruitment, but the most pronounced effect is the loss of GLP-1-induced cAMP generation in the triple AA phospho-null mutant compared to the WT receptor, an effect that results in biased signalling favouring β-arrestin 2 recruitment over cAMP generation and that is significantly mitigated by the inhibition of PDE4 action with rolipram.

## Declaration of interest

The authors have no conflicts of interest to declare.

## Funding

The A.T. lab is funded by grants from the MRC (MR/X021467/1) and the Wellcome Trust (301619/Z/23/Z).

## Acknowledgements

The authors thank the Imperial College Facility for Imaging by Light Microscopy (FILM) for technical support on light microscopy experiments, Dr Jorge Bernardino de la Serna, National Heart and Lung Institute (NHLI), Imperial College London and Dr Affiong I. Oqua, Dept. Metabolism, Digestion and Reproduction, Imperial College London, for support on RICS analysis, and the Section of Endocrinology and Investigative Medicine at Imperial for access to high-content microscope and plate readers.

## Notes

### Competing Interest Statement

The authors have declared no competing interest.

