## Supplementary material for "Glucagon-like peptide-1 receptor C-terminal tail phosphorylation determines signalling responses from pancreatic β-cells": Table 1

| **Primer** | **Sequence** |
| --- | --- |
| S441/2A Forward | 5'-cctcaagtgtcccaccgccgccctgagcagtggagcc-3’ |
| S441/2A Reverse | 5'-ggctccactgctcagggcggcggtgggacacttgagg-3’ |
| S444/5A Forward | 5'-ccaccagcagcctggccgctggagccacggcgg-3' |
| S444/5 Reverse | 5'-ccgccgtggctccagcggccaggctgctggtgg-3’ |
| S451/2A Forward | 5'-ggagccacggcgggcgccgccatgtacacagccac-3’ |
| S451/2 Reverse | 5'-gtggctgtgtacatggcggcgcccgccgtggctcc-3' |
| S444/5A + S441/2A Forward | 5'-caccgccgccctggccgctggagccacggcg-3' |
| S444/5A + S441/2A Reverse | 5'-cgccgtggctccagcggccagggcggcggtg-3’ |
| Sequencing primer | 5'-ccttcacctccttccaggggct-3' |

**Table 1: List of site-directed mutagenesis and sequencing primers used in the study.**
